# Ongoing exposure to peritoneal dialysis fluid alters resident peritoneal macrophage phenotype and activation propensity

**DOI:** 10.1101/2020.03.02.973404

**Authors:** Tara E. Sutherland, Tovah N. Shaw, Rachel Lennon, Sarah E. Herrick, Dominik Rückerl

## Abstract

Peritoneal dialysis (PD) is a more continuous alternative to haemodialysis, for patients with chronic kidney disease, with considerable initial benefits for survival, patient independence and healthcare costs. However, longterm PD is associated with significant pathology, negating the positive effects over haemodialysis. Importantly, peritonitis and activation of macrophages is closely associated with disease progression and treatment failure. However, recent advances in macrophage biology suggest opposite functions for macrophages of different cellular origins. While monocyte-derived macrophages promote disease progression in some models of fibrosis, tissue resident macrophages have rather been associated with protective roles. Thus, we aimed to identify the relative contribution of tissue resident macrophages to PD induced inflammation in mice. Unexpectedly, we found an incremental loss of homeostatic characteristics, anti-inflammatory and efferocytic functionality in peritoneal resident macrophages, accompanied by enhanced inflammatory responses to external stimuli. Moreover, presence of glucose degradation products within the dialysis fluid led to markedly enhanced inflammation and almost complete disappearance of tissue resident cells. Thus, alterations in tissue resident macrophages may render longterm PD patients sensitive to developing peritonitis and consequently fibrosis/sclerosis.

## Introduction

An estimated 5-10 million people worldwide die every year due to chronic kidney disease [1]. In Europe, an average of 850 people per million population (pmp) require renal replacement therapy (RRT) and 120 new patients pmp commence treatment annually [2]. The average 5-year survival rate of patients receiving RRT is only 50.5%. This can be improved to over 90% if patients receive a kidney transplant, but rates of transplantation remain low (32 pmp) primarily due to organ availability, and the majority of patients rely on dialysis as a therapy to substitute excretory kidney function [2].

Peritoneal dialysis (PD), utilizing the body’s own peritoneal membrane as a filter during dialysis, is a cost-effective alternative to haemodialysis (HD). Although PD has been associated with better initial survival rates [3], lower costs for the health system [4] and increased patient autonomy [5] as compared to HD, the incidence rate of PD over several decades has dropped in Europe [3]. There are a variety of reasons for this reduction in PD uptake, but the significant risk of adverse effects and in some cases fatal outcomes, has limited the general adoption of PD in adult patients across Europe [6]. Treatment failure is commonly associated with repeated episodes of peritonitis (i.e. inflammation) and a progressive thickening and vascularisation of the peritoneum, leading to impaired filtration and thus reduced efficacy of PD [6, 7]. In rare cases, the fibrotic changes to the peritoneum become so extreme that they form a fibrous cocoon encapsulating the internal organs, called Encapsulating Peritoneal Sclerosis (EPS), leading to persistent or recurring adhesive bowel obstruction [7-9]. The diagnosis of EPS is an indication to urgently discontinue PD with the mortality approaching 50% within one year after diagnosis [10]. Management of EPS includes surgery but there is a relatively high frequency of symptom recurrence. In contrast, immunosuppressive treatments and the use of anti-fibrotic agents, like tamoxifen, have shown noticeable benefits to patient survival [10]. Therefore, aberrant activation of the immune system appears to be linked to both alteration of peritoneal structure and PD treatment failure, as well as the progression of the fibrotic sequelae. Indeed, experimental rodent models of the disease, have suggested inappropriate or excessive activation of macrophages (MΦ) as a major cause of the pathology [11-14].

Recent advances in MΦ biology have highlighted the intrinsic heterogeneity of MΦ populations [15]. Grossly simplified, MΦ can be split into tissue resident macrophages (MΦres) which are present in tissues during homeostasis, and monocyte-derived macrophages (MΦmono), which are recruited to the tissue during inflammatory conditions [16]. Both MΦ populations can and do respond to external stimuli, like infection, but they possess distinct response profiles and adopt distinct functional properties [17, 18].

In health, peritoneal MΦres are essential for maintaining tissue homeostasis by silently removing apoptotic cells through efferocytosis [19, 20] and by providing a source of tissue-reparative cells that infiltrate surrounding tissues (e.g. the liver) during injury [21]. Importantly, upon encountering inflammatory signals, MΦres undergo the Macrophage Disappearance Reaction (MDR) [22]. Following injection of inflammatory agents (e.g. bacterial antigens) or infection, the number of peritoneal MΦres detectable in peritoneal lavage will drop significantly within a few hours [23]. The exact mechanism underlying MDR is still not completely understood, but it has been proposed that MΦres undergo activation-induced cell death [18] or adhere to the mesothelial lining of the peritoneal cavity reducing their recovery via lavage [21, 24, 25]. Of note, the degree of detectable MΦres -loss directly correlates to the amount of inflammatory stimulus (e.g. cfu of bacteria) and the recruitment of inflammatory MΦmono [18]. Following resolution of inflammation, MΦres can re-populate the peritoneal cavity and return to homeostatic numbers through proliferative expansion of remaining MΦres [26].

In models of fibrotic disorders of the lung or kidney, influx of monocytes and monocyte-derived MΦ has been linked to disease progression and induction of pathology [27-29]. Similarly, in rodent models of peritoneal fibrosis, preventing the influx of monocytes or depleting all MΦ limits the degree of peritoneal thickening and improves glomerular filtration [11, 30, 31]. Moreover, injection of MΦmono, activated ex vivo using bacterial antigens (i.e. lipopolysaccharide), often referred to as M1 MΦ, exacerbates disease progression [14]. Together these data highlight a role for inflammation and infiltration of MΦmono in the progression of peritoneal fibrosis and seem to provide a cohesive picture explaining the enhanced risk of PD failure associated with repeated episodes of peritonitis [32]. In this context, it is interesting to note that continuing peritoneal irrigation and thus continual removal of peritoneal cells, including any inflammatory infiltrate, in patients discontinuing PD, has been suggested to prevent subsequent EPS formation [33].

Other studies in rodents have found a role for anti-inflammatory, IL-4-activated MΦ (also called M2), characterised by the expression of CD206, Arg1 and Ym1, in promoting peritoneal dialysis fluid (PD fluid) induced fibrosis [11, 34, 35]. Additionally, chronic fibrotic kidney disease is linked to a switch from inflammatory M1 to predominantly M2 activated MΦ [36]. Indeed, both types of MΦ activation seem to have the capacity to promote kidney fibrosis, possibly acting during different phases of the pathology [37]. Importantly, some markers used to define M2 are differentially expressed on MΦres and MΦmono. For example CD206 is expressed constitutively by MΦmono but not MΦres [17]. Thus, the described role of M2 MΦ may merely reflect an enhanced influx of MΦmono rather than IL-4-mediated activation. In fact, transfer of exogenously activated M2 MΦ showed no impact on disease progression in a model of peritoneal fibrosis [14] and even reduced pathology in a model of renal fibrosis [38]. Moreover, using different strategies to deplete MΦ in kidney fibrosis yielded opposing results with regard to disease progression indicating that depletion of different subsets of MΦ may lead to different outcomes [39]. Indeed, renal MΦres as compared to MΦmono have been shown to be protective in a model of kidney pathology [29]. This suggests that the cellular origin of MΦ may play a prominent part in determining their role during fibrotic disorders and influence the overall disease outcome.

Here we analysed the effects of repeated PD fluid injection on MΦ population dynamics and responses to activating signals. Our data indicate a significant change in MΦres phenotype over time during PD fluid administration. MΦres lost expression of anti-inflammatory and efferocytic markers correlating with enhanced inflammatory responses to external stimulation. Importantly, the enhanced inflammatory phenotype of MΦres persisted even when PD fluid administration was stopped. Interestingly, Nanostring-transcript analysis revealed reduced expression of genes in the Adenosine / G-protein-receptor coupled pathway indicating a potential cause for the loss of regulatory phenotype in MΦres. In contrast, addition of glucose degradation products, known enhancers of PD-pathology [40], led to strongly enhanced inflammation, virtually complete loss of MΦres and enhanced engagement of the TGF-β-as well as Interferon-associated pathways. Thus, repeated exposure to PD fluid may render patients more susceptible to peritonitis, and by extension, to peritoneal fibrosis due to exaggerated inflammatory responses.

## Results

### Dialysis fluid induces the disappearance of tissue resident MΦ

To determine the impact of peritoneal dialysis fluid (PD fluid) injection on peritoneal MΦ populations (see supplementary Fig S1 for gating strategy) we first characterised the cellular dynamics induced after a single application of PD fluid. Six hours after PD fluid injection MΦres underwent a pronounced disappearance reaction with the number of F4/80 high MΦres reduced to approximately 30% of the levels found in naive control animals (Fig 1A). Simultaneously, a significant influx of neutrophils and Ly6C high monocytes were detected, indicative of an inflammatory response (Fig 1B&C). Of note, similar to previous reports [23] tissue dwelling, monocyte-derived MΦmono (F4/80 low MHC-II high) also underwent a disappearance reaction and were reduced in numbers by approximately 40 % (Fig 1D). F4/80 high MΦres displayed limited signs of activation at this early time point post PD fluid injection. No increased expression of MHC-II could be detected (Fig 1E). In contrast intracellular Ym1 and CD206 expression were significantly enhanced following PD fluid injection, but expression was restricted to less than 10 % of cells (Fig 1F&G). The disappearance of MΦres in this context seemed in part due to enhanced cell death as indicated by increased levels of Annexin V staining (Fig 1H). By 24 hours post PD fluid injection, the numbers of MΦres had returned to baseline levels and expression of CD206 was no longer significantly different, whereas Ym1 expression remained elevated compared to naïve controls (Fig S2).

**Figure 1:**
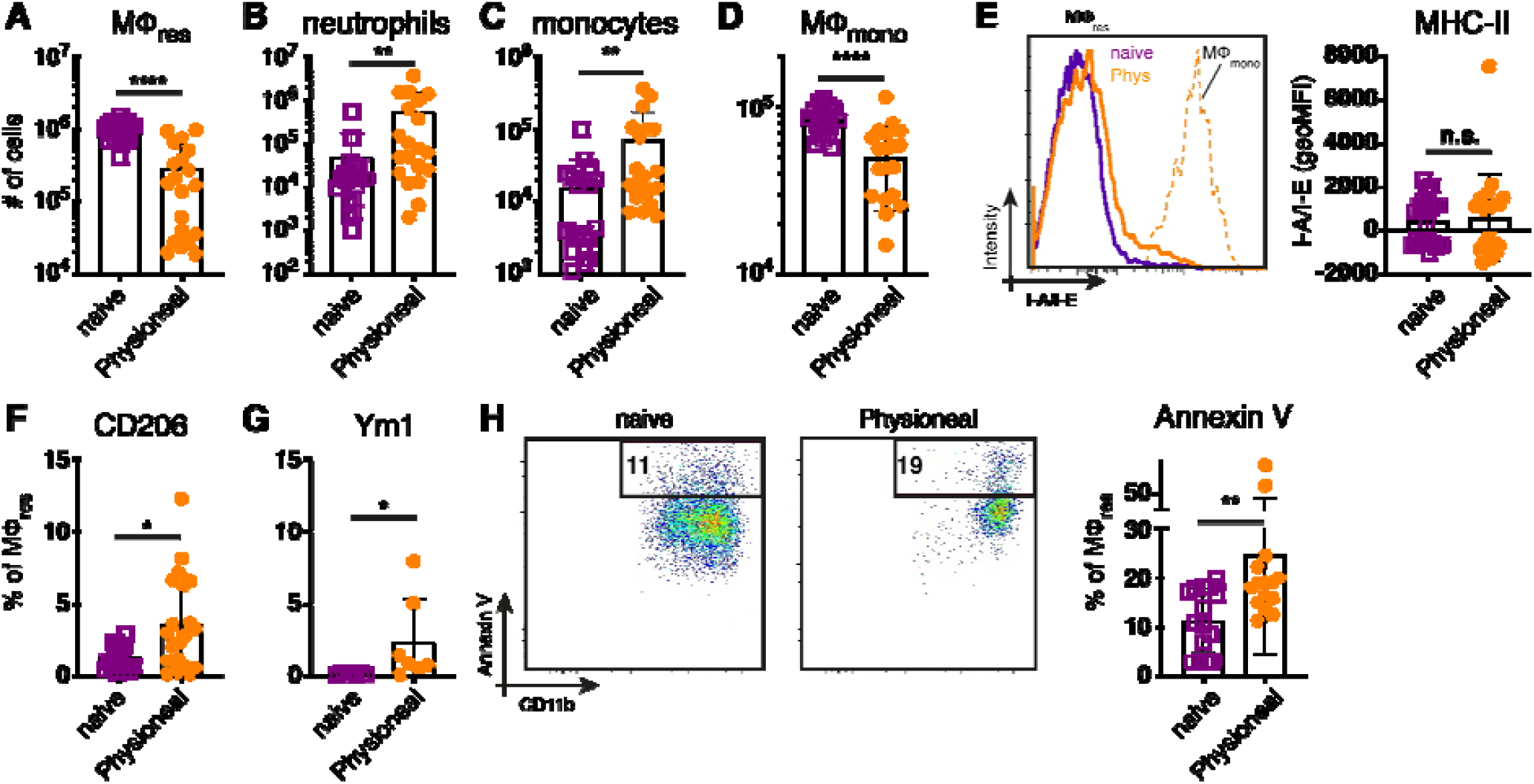
Injection of dialysis fluid alters peritoneal myeloid cell composition. C57BL/6 mice were injected with Physioneal (filled circles) or left untreated (open squares) and whole PEC isolated 6 h later. Cells were counted and analysed by flow cytometry to determine the number of A) MΦres (lineage-CD11b+, F4/80high, MHC-II low), B) neutrophils (lineage +, SSC mid, MHC-II-,F4/80-,CD11b+), C) monocytes (lineage-CD11b+, F4/80-,MHC-II low, Ly6C high) and D) MΦmono (lineage-CD11b+, F4/80low, MHC-II high). E) Histogram and quantitative data of fluorescence mean intensity of MHC-II expression by MΦmono (grey, dashed line) or MΦres from Physioneal injected (black, solid line) or naive animals (grey, solid line). F-H) Expression of CD206, Ym1 and binding of Annexin V by MΦres identified in A) assessed by flow cytometry. Datapoints depict individual animals and bars indicate mean and SD. Data pooled from 4 independent experiments using 3-5 animals per group. Data analysed using a Mann-Whitney-U test after transformation. n.s.: not significant; *: p<0.05; **: p<0.01; ****: p<0.0001 lineage: TCRβ CD19 Siglec-F Ly6G

Overall, these data are consistent with the induction of low grade inflammation caused by PD fluid injection accompanied by a significant change in the prevalence of various myeloid cell populations within the peritoneal cavity.

### Repeated PD fluid treatment induces a gradual change in MΦres phenotype

We next sought to determine the long term effect of PD fluid administration on peritoneal MΦ populations. For this, animals were injected once per day, 5 times a week with PD fluid i.p. and the peritoneal exudate cells (PEC) analysed 24 hours after 1, 4, 9 and 14 injections, respectively (Fig 2A). Over the course of the experiment the number of MΦres remained consistently lower in Physioneal treated animals as compared to naive mice (Fig 2B). Simultaneously, enhanced influx of Ly6C monocytes and successive accumulation of F4/80 low MΦmono could be detected in the peritoneal cavity following multiple rounds of PD fluid injection (Fig 2C& D).

**Figure 2:**
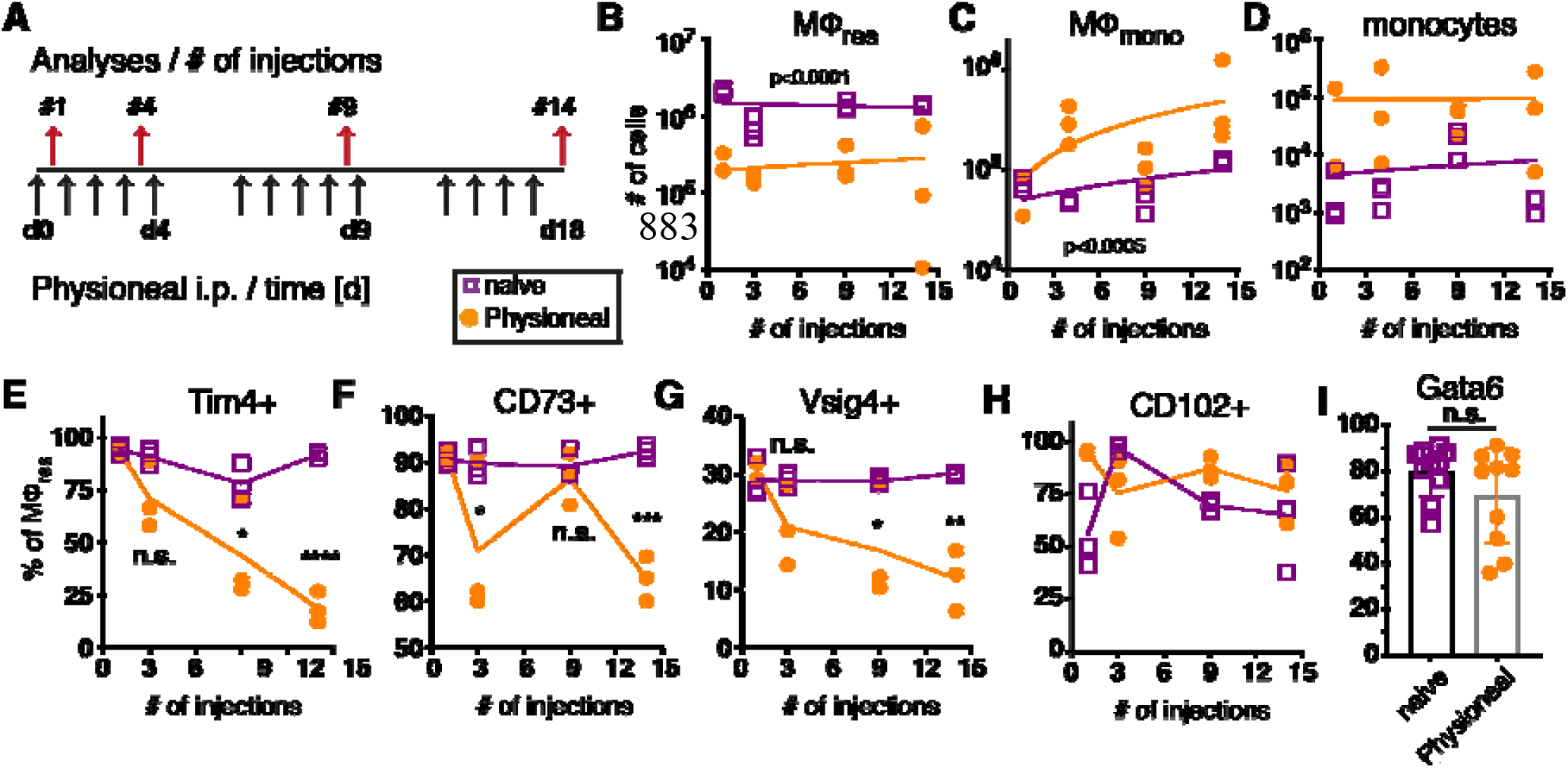
Repeated injection of PD fluid leads to a progressive change in MPres phenotype. C57BL/6 mice were injected with PD fluid (filled circle) or left untreated (open square) 5 times a week for the indicated number of injections. At each timepoint whole PEC were isolated 24 h after the last injection and analysed by flow cytometry. A) Schematic depiction of experimental timeline. B-D) Number of MΦres, MΦmono and Ly6C high monocytes. E-H) Expression of MΦres associated cell-surface markers on F4/80 high MΦ. I) Intracellular expression of Gata6 in MΦres identified in A). Datapoints depict individual animals and lines indicate (B-H) median or (I) mean with SD. Data from a single experiment using 3 animals per group per timepoint. Data analysed using 2-way ANOVA followed by Tukey’s post-hoc test after transformation. n.s.: not significant; *: p<0.05; **: p<0.01; ****: p<0.0001

Despite the overall reduced numbers of MΦres, repeated injection of PD fluid led to marked induction of Ki67 expression in MΦres, indicative of proliferative expansion (Fig. S3). This is in line with a previous report demonstrating repopulation of the peritoneal cavity by MΦres following an inflammatory insult through proliferation [41]. Thus, it is likely that PD fluid injection leads to a repeated cycle of MΦres disappearance followed by repopulation. Similarly, it is likely that the gradual influx of monocytes and MΦmono underlies cyclic fluctuations after each round of PD fluid injection. Thus, this data indicated that MΦ populations within the peritoneal cavity undergo dynamic but limited changes following PD fluid instillation, with a slight accumulation of MΦmono over time.

Closer investigation of the MΦres phenotype, however, revealed a progressive loss of characteristics that define the tissue resident phenotype. In particular markers associated with the efferocytic and anti-inflammatory function of MΦres were expressed at increasingly lower levels following repeated injection of PD fluid (Fig 2E-G). T cell immunoglobulin and mucin domain containing 4 (Tim4), a molecule associated with the efficient removal of apoptotic cells [42, 43] as well as V-set and immunoglobulin domain–containing 4 (Vsig4), associated with limiting inflammatory responses [44, 45], are specifically expressed in MΦres during steady state conditions [46]. MΦres gradually lost expression of these markers with increasing number of PD fluid injections (Fig 2 E & G). Similarly, CD73, an anti-inflammatory effector molecule specifically expressed by MΦres [47, 48] was found to be significantly reduced after 14 injections of PD fluid (Fig 2 F).

In contrast, expression of CD102 (ICAM2) was not altered by PD fluid injection (Fig 2 H). A similar phenotype (loss of MΦres marker expression with sustained CD102 expression) has previously been described in mice lacking the transcription factor Gata6, indicating that repeated PD fluid injection may exert its effects via affecting Gata6 expression [49]. However, no significant difference in Gata6 expression by MΦres was found after 9 injections of PD fluid measured in subsequent experiments (Fig 2 I). Notably, Gata6 expression levels varied following PD fluid injection with diminished expression detected in approximately 50 % of animals (Fig 2 I). Thus, any effect of PD fluid injection on Gata6 expression may be transient or reflect the impact of other factors, like the degree of inflammation.

Taken together, following repeated PD fluid injection, MΦres, while retaining their tissue identity (F4/80 high CD102+ Gata6+), loose some of their functional characteristics and in particular anti-inflammatory and efferocytosis associated functions.

### Repeated PBS injection but not chronic bacterial infection leads to a loss of MΦres characteristics

As shown in Figure 1 and previously reported [14], repeated injection of PD fluid is associated with the induction of low grade inflammation as evidenced by influx of neutrophils and Ly6C+ monocytes. Thus, to assess whether the loss of MΦres functional markers (i.e. Tim4, CD73, Vsig4) was due to the elicited inflammatory response, we analysed peritoneal exudate cells from animals subjected to prolonged bacterial infection. Animals were infected orally with attenuated Salmonella enterica ser. Typhimurium (SL3261, ΔaroA) and peritoneal cells analysed during the chronic phase of infection, 34 and 55 days post infection. In line with previous data [18] and confirming the inflammatory environment, MΦres from SL3261 infected animals showed clear upregulation of MHC-II as well as enhanced expression of Sca-1 on day 55 (Fig 3A & B). Moreover, persistent influx of neutrophils was detected in the peritoneal cavity of SL3261 infected animals (Fig 3C) as well as live, cfu-forming bacteria (Fig 3D) confirming an ongoing pro-inflammatory activation.

**Figure 3:**
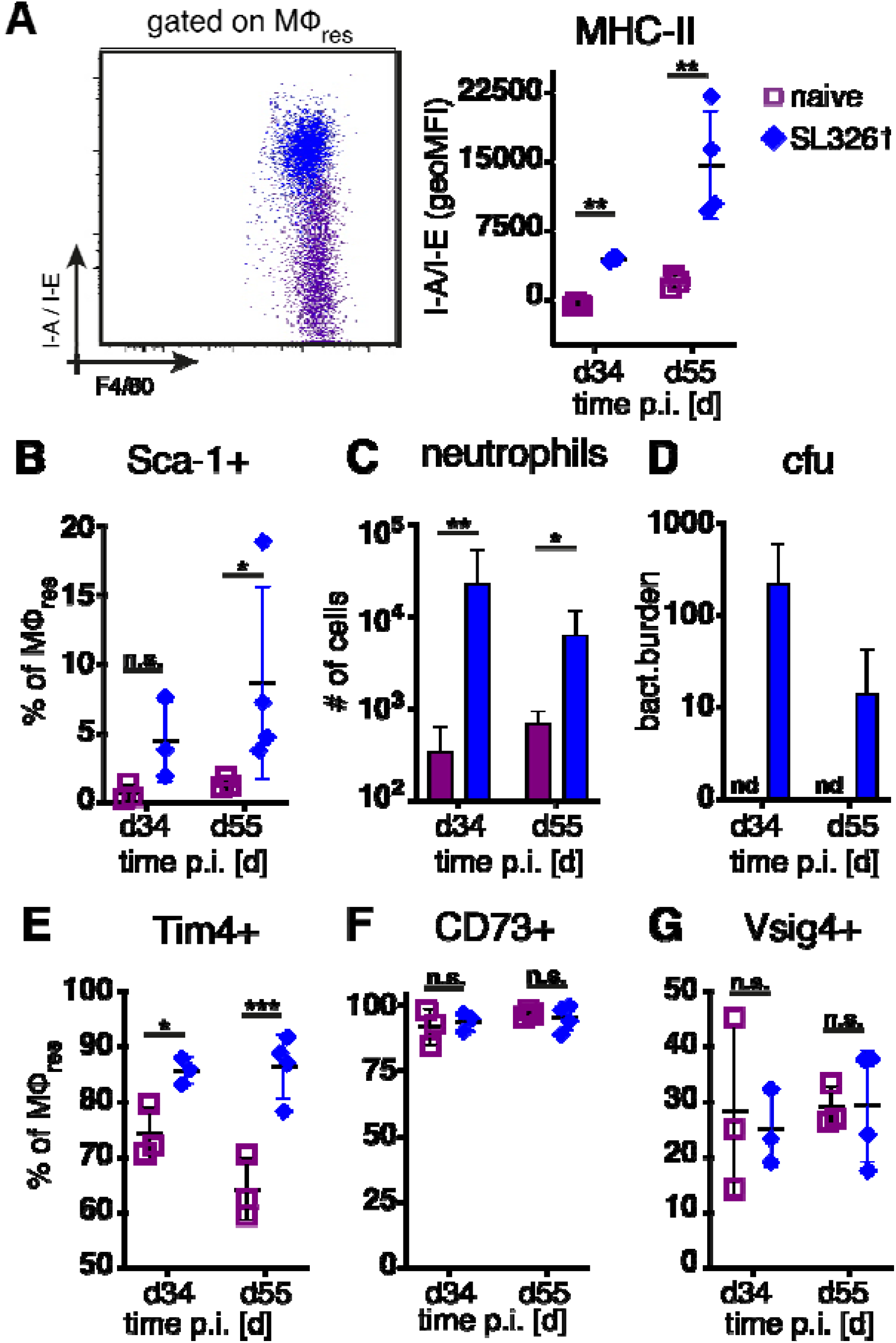
Chronic inflammation induced by S.Typhimurium infection does not phenocopy repeated PD fluid injection. C57BL/6 mice were infected orally with 1×10^8 cfu S.Typhimurium (closed rhombus) or left untreated (open squares) and peritoneal cells analysed 34 (d34) and 55 days (d55) post infection. Dotplot (d55) and quantification of MHC-II (A) and Sca-1 (B) expression by MΦres assessed by flow cytometry C) Number of peritoneal neutrophils. D) Enumeration of bacterial colony forming units present in the peritoneal cavity. E-G) Expression of MΦres associated cell surface markers assessed by flow cytometry. Datapoints depict individual animals and bars / lines indicate mean with SD. Data representative of one (d34) or two (d55) separate experiments using 3-4 animals per group per timepoint. Data analysed using 2-way ANOVA followed by Tukey’s post-hoc test after transformation. n.s.: not significant; *: p<0.05; **: p<0.01; ***: p<0.001

However, unlike following repeated PD fluid injection, MΦres isolated from animals harbouring S. Typhimurium did not show any noticeable loss of Tim4, CD73 or Vsig4 expression in the chronic phase of the infection (Fig 3 E-G). Rather to the contrary MΦres from SL3261 infected animals expressed elevated levels of Tim4 (Fig 3 E). Thus, chronic inflammatory conditions alone did not cause the progressive loss of MΦres phenotype as observed following repeated Physioneal injection.Importantly, injection of sterile PBS instead of PD fluid induced similar, albeit less pronounced changes in peritoneal MΦ phenotype (Fig 4A). This would indicate that repeated disturbance of the peritoneal immune system alone, rather than specific constituents of the dialysis fluid, was sufficient to drive the observed alterations in peritoneal MΦres. However, PD fluid enhanced these effects (Fig 4A).

**Figure 4:**
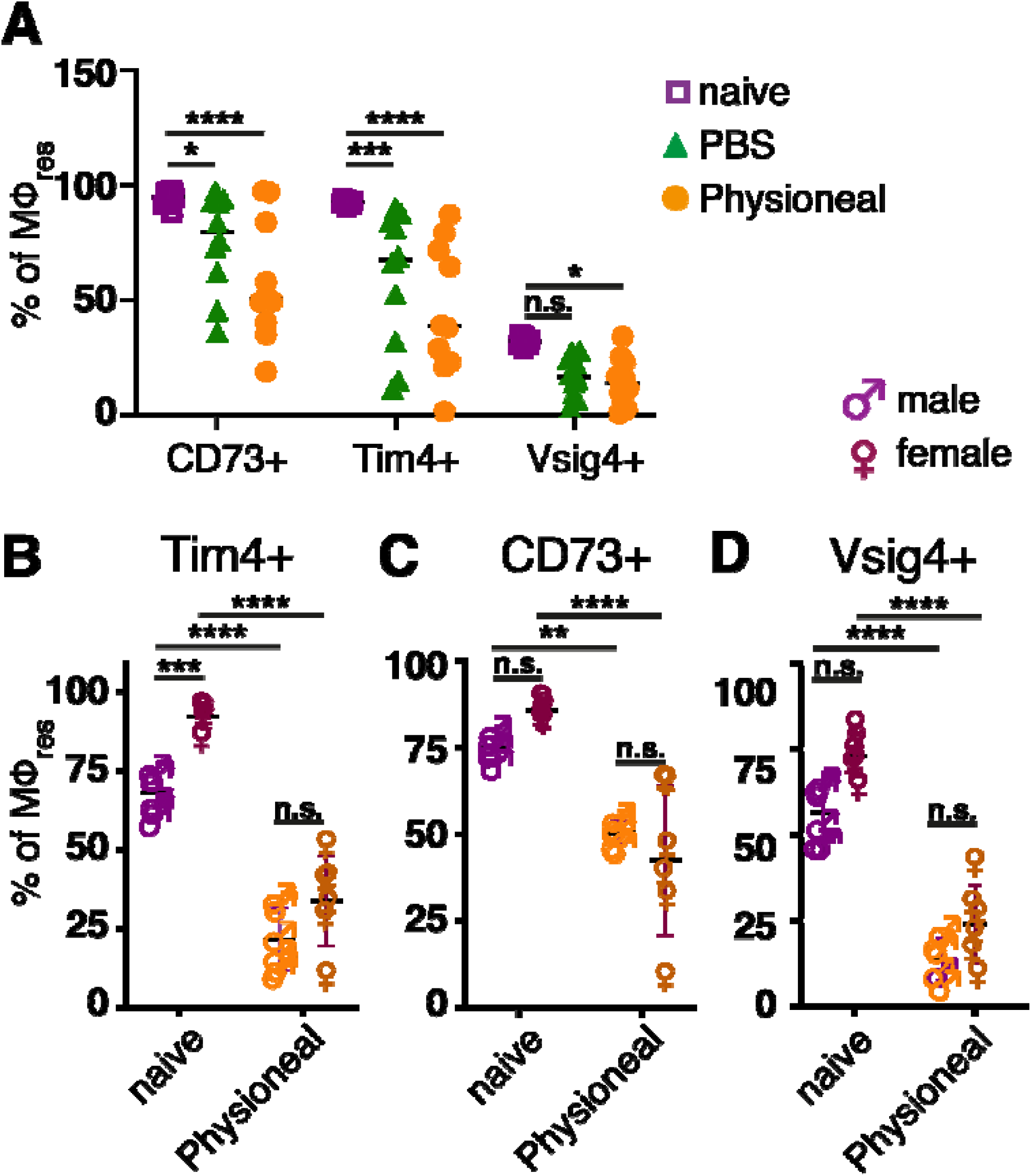
Repeated injection of sterile saline induces similar alterations to dialysis fluid. A)Female C57BL/6 mice were injected 5 times a week for a total of nine injections with Physioneal (circles), sterile PBS (triangles) or left untreated (squares). 24 h after the last injection whole PEC were isolated and analysed by flow cytometry for expression of MΦres associated cell-surface markers. Data pooled from two independent experiments. B)Male (♂) and female (♀) C57BL/6 mice were injected 5 times a week for a total of nine injections with PD fluid (dark symbols) or left untreated (light symbols). 24 h after the last injection whole PEC were isolated and analysed as described for A). Datapoints depict individual animals and lines indicate mean and SD. Data from a single experiment. Data analysed using 2-way ANOVA followed by Tukey’s multiple comparison test after transformation. n.s.: not significant; *: p<0.05; **: p<0.01; ****: p<0.0001

Furthermore, it has been suggested that male peritoneal dialysis patients have a significantly reduced survival rate on PD than female patients [50]. Although the reason for this discrepancy is unknown and likely due to multiple factors, differences in the inflammatory response may contribute to the observed effects. Moreover, male and female mice differ considerably in the maintenance and cellular dynamics of peritoneal MΦ [51]. Thus, we assessed the effect of repeated PD fluid injection on myeloid cell populations in male and female mice. MΦres from naive male or female mice showed comparable expression levels of CD73 and Vsig4. In contrast naive male mice showed considerably lower expression of Tim4 (Fig 4B-D). However, independent of these differences in the steady state, Physioneal induced the loss of Tim4, CD73 and Vsig4 in both sexes to a similar degree. Taken together this data shows that repeated disturbance of the peritoneal environment triggers low grade inflammation which alters the MΦres phenotype.

### Prolonged PD fluid injection alters the response of MΦres to external stimuli

Our previous data had highlighted a significant loss of Tim4 as well as Vsig4 and CD73, markers which have been associated with MΦres core functions (i.e. efferocytosis and anti-inflammatory activity) [42, 45, 52]. Thus, we next aimed to assess whether repeated injection of dialysis fluid altered MΦ functional responses.

To test the capacity of MΦres to take up and ingest apoptotic cells, whole PEC from animals injected for various times with Physioneal or from naive controls were incubated in the presence of apoptotic thymocytes labelled with pHrodo. pHrodo labelled cells emit a very low, nearly undetectable fluorescent signal after staining, but will become brightly fluorescent and clearly detectable by flow cytometry after encountering an acidic environment, as found inside a phagolysosome [53]. Thus, use of pHrodo allows the reliable detection of ingested apoptotic cells as compared to labeled cells bound to the surface of a phagocyte.

Repeated injection of Physioneal gradually reduced the proportion of myeloid cells (total CD11b+lin-) capable of ingesting apoptotic cells (Fig 5A). This was in part due to the increased proportion of Ly6C high monocytes and F4/80 low, monocyte-derived MΦ within the myeloid cell pool (Fig 2C&D), cells which possess reduced efferocytic activity [54]. However, when the analysis was restricted to MΦres a similar progressively reduced capacity to ingest apoptotic cells was detected (Fig 5B). Thus, MΦres from Physioneal injected animals seemed to lose the functionality to carry out efferocytosis efficiently. Of note, this loss of efferocytic capacity was not restricted to the use of dialysis fluid, as repeated injection of sterile PBS induced a similar reduction in the capacity of MΦres to ingest apoptotic cells (Fig S4A).

**Figure 5:**
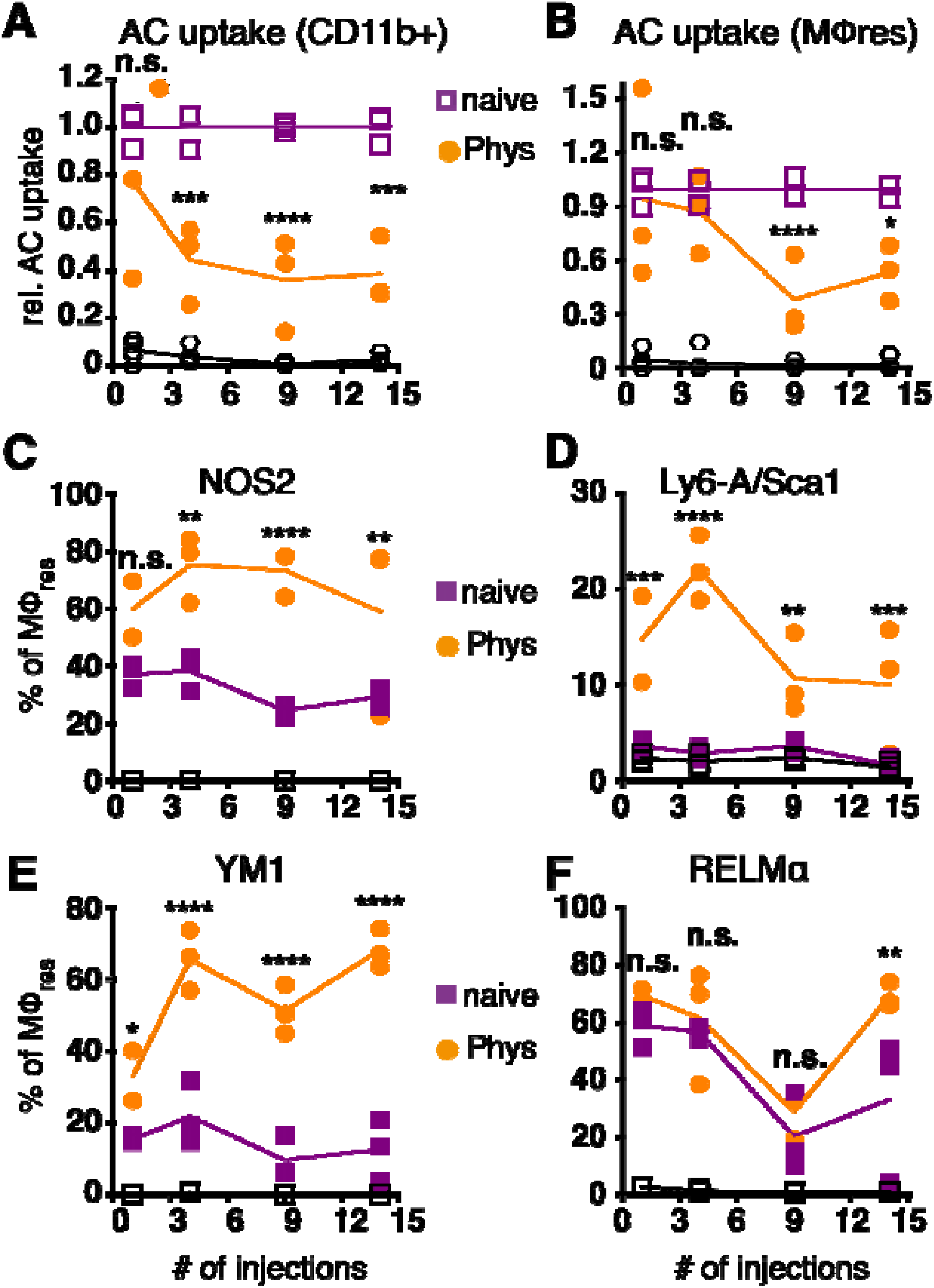
MΦres from PD fluid-injected animals show progressively enhanced responses to stimulation. C57BL/6 mice were injected with PD fluid (circles) or left untreated (squares) 5 times a week for the indicated number of injections. At each timepoint whole PEC were isolated 24 h after the last injection and incubated in vitro in the presence of pHrodo labelled apoptotic thymocytes for 90 minutes (A & B) or stimulated with LPS/IFNγ (C & D; 6 h) or recombinant IL-4 (E & F; 24 h). Uptake of apoptotic cells by total CD11b+ myeloid cells (A) or MΦres (CD102+I-A/I-E low) (B) as well as expression of MΦres associated cell-surface markers (C-F) assessed by flow cytometry. Datapoints depict individual animals and lines indicate mean and SD. Data from a single experiment. Data analysed using 2-way ANOVA followed by Tukey’s post-hoc test after transformation. n.s.: not significant; *: p<0.05; **: p<0.01; ****: p<0.0001

To further analyse whether MΦres from Physioneal injected animals were in general less responsive to external stimuli, we subjected whole PEC to in vitro stimulation with LPS and rIFNγ. MΦres from PD fluid treated animals showed a gradually increasing inflammatory response towards bacterial stimulation (Fig 5C&D). The proportion of cells expressing NOS2 or Sca-1, markers of pro-inflammatory M1 activation [55, 56], was consistently higher in cells from PD fluid treated animals as compared to cells from naive animals (Fig 5C&D). Indeed, Sca-1 was not found to be upregulated on naive MΦres after 6 h stimulation with IFNγ/LPS in vitro, indicating repeated Physioneal injections resulted in a stronger and more rapid response of MΦres to pro-inflammatory stimuli.

Importantly, an enhanced activation profile was not only observed in response to pro-inflammatory stimuli, but also in response to rIL-4, a driver of M2, anti-inflammatory MΦ activation [57]. MΦres stimulated with rIL-4 for 24 h showed increased expression of Ym1 and at a later timepoint also Relmα (Fig 5 E&F).

Thus, MΦres altered their functional repertoire following repeated exposure to dialysis fluid, with reduced efferocytic capacity and enhanced responses to both M1 and M2 polarising stimuli.

### Altered PD fluid-induced responsiveness of MΦres is maintained after treatment is discontinued

Next, we examined whether the phenotypical and functional changes observed in MΦres following repeated PD fluid injection were temporary or persisted even after treatment ceased. For this, mice were injected 5 times a week with Physioneal or PBS for a total of 9 injections, a timepoint when the altered MΦres phenotype is evident (Fig 2), and then rested for seven days. Subsequently, the PEC were collected and analysed for the expression of activation markers as well as re-stimulated in vitro to analyse their response to external stimuli as described above. MΦres from discontinued Physioneal or PBS injected animals maintained a reduced expression of CD73, Tim4 and Vsig4 compared to naive controls (Fig 6A). Similarly, MΦres from Physioneal and PBS injected animals showed an enhanced response to in vitro stimulation with LPS/IFNγ (NOS2, Sca1; stimulated for 6 h) (Fig 6B). To verify whether the altered MΦ phenotype observed in rested MΦres was due to the integration of inflammatory, monocyte-derived cells into the resident pool we conducted lineage tracing experiments using Cx3cr1^CreER^:R26-eyfp mice [58]. Animals were dosed daily during the first 5 days of the experiment with Tamoxifen by oral gavage. This method will efficiently label peritoneal monocyte-derived MΦ populations (eg. MΦmono) while MΦres remain unlabelled. This allows to determine the degree of integration of monocyte-derived cells into a resident MΦ population [58]. In addition, all animals were treated as described above (9 injections of Physioneal, 5 times per week) and rested for a further seven days (Fig 6C). In our hands approximately 20 % of Ly6C-CD115+ blood monocytes stained positive for eYFP two weeks after the last tamoxifen administration confirming efficient labelling. Moreover, the degree of labelling found in blood monocytes was independent of any treatment with Physioneal or PBS (suppl.Fig S5). In contrast, approximately 2.5 % of peritoneal F4/80 high MΦres from naive animals were eYFP+ (Fig 6D) confirming their maintenance is partly independent of monocytic influx [59]. Animals injected with Physioneal exhibited a slightly higher proportion (∼ 5 %) of eYFP+ MΦres (Fig 6D), indicating enhanced integration of monocyte derived cells into the resident pool. However, independent of any treatment, the proportion of eYFP+ cells in MΦres did not reach the same levels as observed in F4/80 low MHC-II high MΦmono (∼ 12%; Fig 6D). Thus, although Physioneal treatment did increase the rate at which monocyte-derived cells integrated into the MΦres pool, the population remained to a significant part resident in origin. Of note, the enhancement of MΦres turnover and monocyte-integration was similarly observed in PBS treated animals (Fig 6D). Importantly, MΦres from tamoxifen-treated Cx3cr1^CreER^:R26-eyfp mice showed similar behaviour in response to repeated Physioneal / PBS injection as observed above, as indicated by a loss of CD73, Tim4 and Vsig4 expression (suppl. Fig S5). Therefore the observed changes in MΦres phenotype can not be explained due to enhanced integration of monocyte derived MΦ into the resident pool alone.

**Figure 6:**
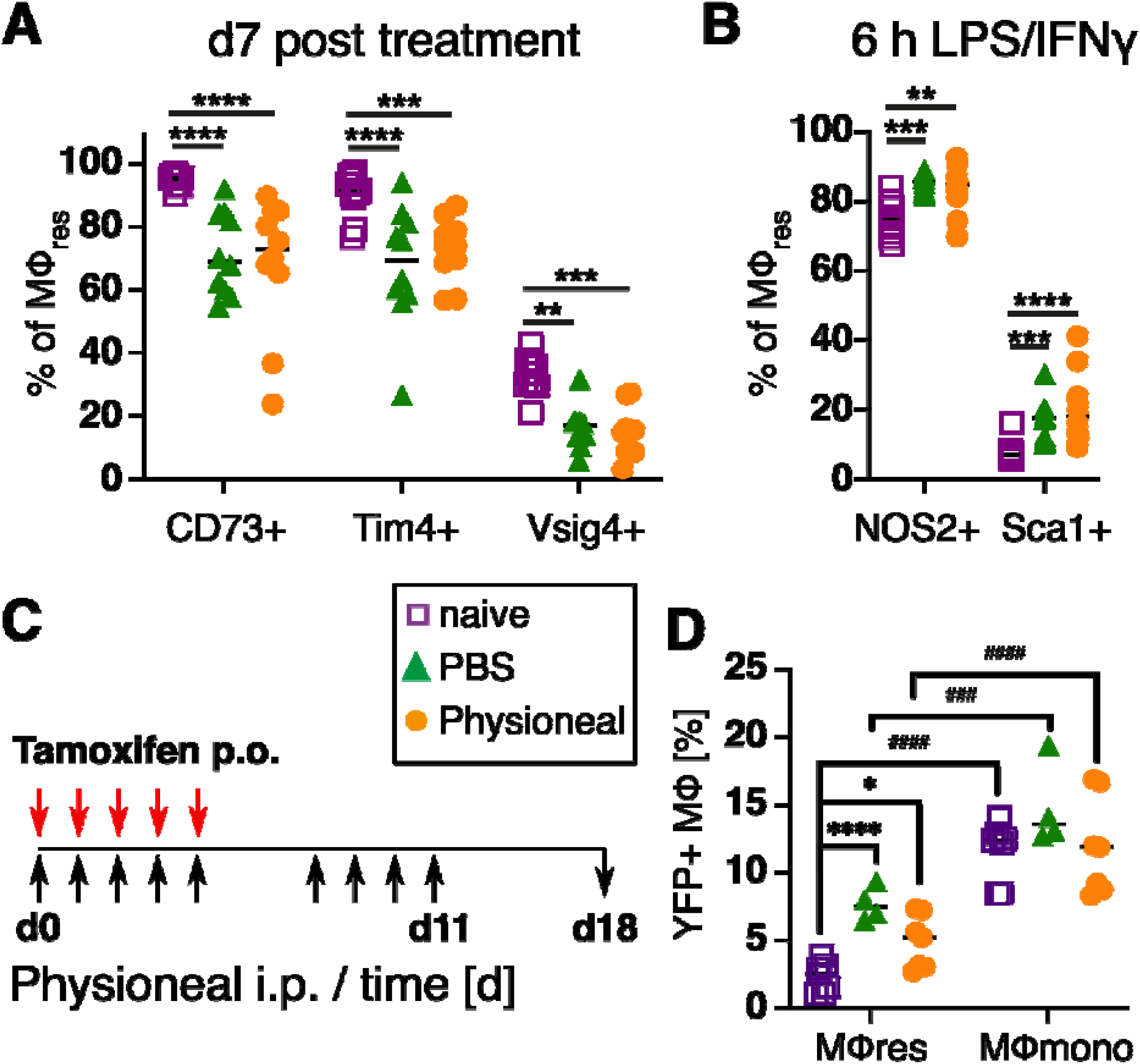
PD fluid conditioned MΦres maintain altered activation phenotype after treatment is discontinued while remaining largely tissue-resident derived. (A) C57BL/6 mice were injected 5 times a week for a total of nine injections with Physioneal (circles), sterile PBS (triangles) or left untreated (squares). 7 days after the last injection whole PEC were isolated and analysed by flow cytometry for expression of MΦres associated cell-surface markers. (B) The cells isolated in A) were subjected to in vitro stimulation with LPS/IFNγ (6 h) and analysed by flow cytometry for NOS2 & Sca1 expression. C&D) To determine the cellular origin of the MΦres Cx3cr1CreER:R26-eyfp mice were treated as described in A) and additionally received 5 doses of tamoxifen per oral gavage at the beginning of the experiment. (C) Schematic depiction of the experimental timeline for experiments with tamoxifen injection. (D) Analysis of eYFP expression by MΦres and MΦmono in Cx3cr1^CreER^:R26-eyfp mice treated with PBS, Physioneal or untreated. Datapoints depict individual animals and lines indicate mean & SD. (A, B) Data pooled from 2 independent experiments. Data analysed using 2-way ANOVA followed by Tukey’s multiple comparison test after transformation. (D) Data from 2 independent experiments analysed by 2-way ANOVA followed by Tukey’s multiple comparison test after transformation. # indicate statistical differences between MΦres and MΦmono of the same treatment. n.s.: not significant; *: p<0.05; **: p<0.01; ***: p<0.001; ****: p<0.0001;

These data highlight that repeated PD fluid injections sensitise resident peritoneal MΦ and, thus, potentially exacerbate the severity and the detrimental sequelae of peritonitis events.

### Glucose degradation products strongly enhance inflammatory responses and loss of tissue resident MΦ markers

Since our previous data revealed limited differences between injection of dialysis fluid and injection of sterile saline solution (PBS), we wanted to verify whether addition of glucose degradation products (GDP), altered the observed MΦ responses. GDP can form in dialysis fluid due to the heat sterilisation process [60] and have been previously shown to strongly enhance PD-associated fibrosis in humans and mice [40, 61], Thus, animals were injected with Physioneal containing 40 mM Methylglyoxal (MGO) for 14 injections and analysed 24 h after the last injection. Addition of MGO to the dialysis fluid dramatically altered the immune profile of the mice, leading to strongly enhanced inflammatory influx, as indicated by significantly enhanced numbers of neutrophils, Ly6C+ monocytes and F4/80 low MHC-high MΦmono (Fig 7A). Moreover, the loss of Tim4, CD73 and Vsig4 expression in F4/80 high (CD11b+ lineage-) cells observed following Physioneal injection was even further enhanced by MGO supplementation (Fig 7 C). However, although the number of cells within the F4/80 high (CD11b+ lineage -) gate were not altered by the addition of MGO (Fig 7 A), these were likely not tissue-resident derived MΦ as indicated by their altered flow-cytometry profile (Fig 7B), as well as the loss of CD102, a marker of tissue residency in peritoneal MΦ [49], which remained unaffected by injection of Physioneal alone (Fig 7C). In addition, Physioneal supplemented with MGO induced significant inflammatory activation of total peritoneal myeloid cells (CD11b+ lineage-cells) as indicated by enhanced expression of Sca-1 and NOS2 (Fig 7D). Furthermore, MΦ markers associated with fibrosis, CD206 and Ym1, where exclusively upregulated in MGO-treated, highly inflammatory settings. In contrast arginase 1 (Arg1) was upregulated in Physioneal treated animals, but not in Physioneal + MGO treated animals (Fig 7 D). Thus, the presence of glucose degradation products strongly enhanced the inflammatory response leading to an almost complete disappearance of homeostatic MΦres.

**Figure 7:**
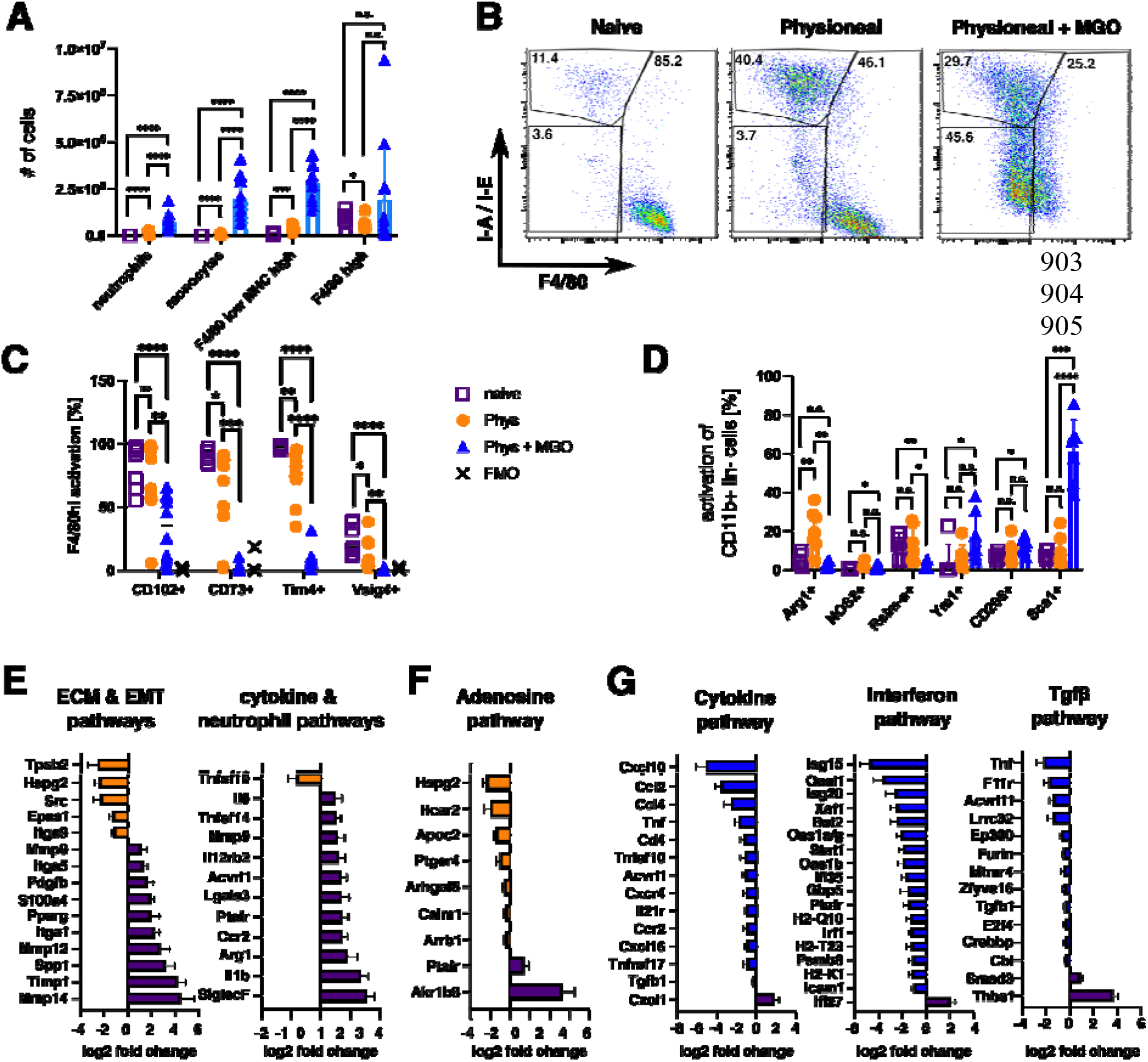
Addition of glucose-degradation products strongly enhances peritoneal inflammation. C57BL/6 mice were injected with PD fluid (Phys, circles) or PD fluid supplemented with 40 mM Methylglyoxal (MGO, triangles) or were left untreated (naive, squares) 5 times a week for a total of 14 injections and analysed 24 h after the last injection. (A) Number of inflammatory cells isolated from the peritoneal cavity. (B) Sample dot plots depicting myeloid cells (CD11b+, lineage -) isolated from the animals in A). (C &D) Expression of MPres associated surface markers (C) as well as MΦ activation markers (D). Datapoints depict individual animals and bars indicate mean and SD. Data from 3 independent experiments. Data analysed using 2-way ANOVA followed by Tukey’s post-hoc test after transformation. n.s.: not significant; *: p<0.05; **: p<0.01; ***: p<0.001; ****: p<0.0001 (E-G) nCounter fibrosis panel analysis of whole PEC isolated in A). Bars indicate genes upregulated in naive (purple), Physioneal (orange) or Physioneal + Methylglyoxal (blue) treated animals. Data depicted as log2 fold change of significantly differentially expressed genes (p<0.05) as mean and SEM of three animals per group.

To further elucidate the differences between PD-fluid alone and PD-fluid enriched with glucose degradation products, we analysed gene expression of total peritoneal exudate cells using the Nanostring nCounter Fibrosis panel platform. Due to legal animal welfare restrictions on the maximum number of i.p. injections we were allowed to give in our experiments, we were unable to detect any histological signs of fibrosis in our model (data not shown). However, in line with previous publications demonstrating considerable thickening of the peritoneum within 5 -8 weeks of daily PD-fluid instillation [11, 61], injection of Physioneal led to the significant up regulation of a series of genes associated with ECM-remodelling and epithelial-mesenchymal transition (EMT) (Fig 7 E). Moreover, in line with the low level inflammation detected by flow cytometry, genes associated with cytokine & neutrophil pathways were significantly enhanced in Physioneal treated animals. Of note, genes associated with the Adenosine-& G-protein coupled receptor signalling pathway were significantly reduced in Physioneal treated animals (Fig 7 F) in line with our finding of reduced CD73 expression (Fig 7 C). Moreover, addition of MGO led to a further increase in expression of inflammatory genes and in particular Interferon-related genes (Cytokine & Interferon pathways; Fig 7 G) in accordance with the enhanced influx of inflammatory cells (Fig 7 A). Furthermore, co-administration of MGO also led to marked increases in expression of Tgf-β-pathway associated genes (Fig 7 G), in line with the previously described enhancement of fibrosis induced by glucose degradation products [61].

Taken together these data highlight, that the presence of glucose degradation products dramatically alters the immune profile within the peritoneal cavity and that distinct gene signatures are associated with steady PD-fluid instillation as compared to pathological conditions.

## Discussion

MΦ are versatile cells implicated in many pathologies [62]. They play both essential protective as well as detrimental / pathological roles, often within the same disease setting [63-65]. Thus, the regulatory mechanisms governing MΦ responses have become the focus of current research aiming to dissect these contradictory behaviours. Moreover, scientists have proposed MΦ as an excellent target for therapeutic interventions, as altering their phenotype may not only improve disease outcome, but actively revert associated pathologies. Here we show, that tissue resident, peritoneal MΦ gradually lose their homeostatic, anti-inflammatory phenotype following PD-fluid instillation and, in particular, a loss of regulation via the Adenosine pathway. In addition we show that the presence of glucose degradation products, known enhancers of PD-associated pathology [40], induces strong inflammatory responses characterised by enhanced Tgf-β-and Interferon signalling.

Recent discoveries have highlighted the inherent diversity of the MΦ pool, indicating the presence of different types of MΦ with different ontogeny [59], response profiles [17] and functional roles [18]. In particular, MΦres and acutely recruited MΦmono have been identified as distinct mediators of pathology. During lung fibrosis recruited MΦmono have been shown to be essential drivers of the fibrotic pathology and disease progression [27]. Similarly, in murine models of peritoneal dialysis associated fibrosis, infiltration of MΦmono has been shown to be detrimental to disease outcome [11, 14, 30, 31]. Furthermore, significant changes in the prevalence of specific MΦ/monocyte populations have been described in dialysis patients, dependent on the history of peritonitis episodes [12]. Thus, the pathology inducing effect of peritonitis associated inflammation in peritoneal dialysis seem to be due to the recruitment and accumulation of fibrosis promoting cells like MΦmono. Similar to these findings, we observed low-grade inflammation and infiltration of MΦmono in our system as well as strongly enhanced influx of MΦmono upon injection of MGO, a driver of PD-associated pathology [61]. However, in addition, we demonstrated a drastic change in phenotype of MΦres. Importantly, this change in phenotype translated into an enhanced response profile boosting pro-inflammatory effector molecule production, which was maintained even when PD-fluid instillation was stopped. Thus, the increased risk of developing pathological sequelae following repeated episodes of peritonitis in PD patients is likely due to the damaging effect of the inflammatory response [66]. But our data implies that patients on longterm PD treatment are at an increased risk to develop clinical symptoms of peritonitis due to an enhanced inflammatory response of peritoneal resident cells. Intriguingly, in PD patients receiving oral supplementation with vitamin D, a factor involved in the expansion of peritoneal MΦres in mice [67], antibacterial responses were enhanced [68]. However, long-term consequences of this enhanced inflammation have not been investigated.

On the other hand, we have found enhanced expression of Arg-1, particularly in Physioneal treated animals. Although, Arg-1 is typically considered a pro-fibrotic marker [69], expression of Arg-1 specifically by macrophages has been shown to limit excessive ECM deposition in models of liver fibrosis [70] and animals lacking Arg-1 expression specifically in myeloid cells fail to resolve atherosclerotic inflammation and disease progression [71]. Taken together with the fact that pathology-enhancing factors like peritonitis will lead to the disappearance of MΦres, as also visible in our experiments using MGO, hints at a potential protective / anti-fibrotic role of MΦres, as has previously been described in the kidney [39] and lung [72, 73]. Indeed, targeting macrophages to induce an anti-fibrotic phenotype has previously been suggested [69]. Moreover, recent single cell analysis of pro-resolving and pro-fibrotic macrophages following injury revealed the presence of a unique macrophage population with enhanced phagocytosis and a gene expression signature closely resembling resident macrophages [74, 75]. Thus, it may be feasible to develop therapeutic approaches fostering a low-inflammatory, pro-resolving environment by targeting and promoting homeostatic MΦres. Our data from Salmonella enterica ser. Typhimurium infected mice indicated that the late stages of infection were linked to an increase in Tim4 expressing MΦres as compared to naive animals. Similarly, previously published results from helminth infected animals indicate enhanced expression of Tim4 on MΦres during the chronic phase of these infection [76]. Hence, factors associated with the resolution of inflammation or chronic Th2 immune responses may allow to ’rejuvenate’ MΦres to regain some of their homeostatic / anti-inflammatory activities. Further research is required to determine the specific factors driving the MΦres phenotype as well as their potential use as therapeutic adjunct during PD.

Independent of the therapeutic potential discussed above, analysing the immune cells contained in the effluent of PD patients may yield a useful biomarker strategy. Changes in the composition of myeloid cells as described by Liao et al. [12] as well as the assessment of cellular activation markers, in particular their Interferon signature, as described in mice here, may allow risk-based patient stratification. Patients likely to develop pathological sequelae of PD treatment could then be prioritised for kidney transplant or transferred to haemodialysis prior to PD failure or before overt pathology develops.

Lastly, our data comparing PBS injected animals to Physioneal injection indicated a limited impact of the composition of the PD fluid on the observed physiological changes. In contrast, our data using MGO as well as previously published reports have identified several constituents of PD fluid, in particular lactate, GDP and low pH, to be instrumental in driving PD related pathology [77, 78]. However, while these factors have a clear impact on the inflammatory and pathological response, our data is in line with a recent report investigating the use of so-called biocompatible PD fluid in patients. Despite considerably lower levels of GDP and lactate in these solutions, the morphological changes observed were very similar to those seen in patients utilising standard PD fluid and even an early increase in microvascular density was detected [79]. Of note, it has been suggested that this adverse effect of biocompatible PD fluid may be due to an altered inflammatory response [80]. Thus, together with our data this suggests it may be more effective to prevent pathological changes by targeting the elicited immune response to PD fluid instillation, even in the absence of peritonitis. Due to this we also chose naive animals as comparator, as these represent healthy patients not currently on PD, and hence the situation any therapeutic intervention should aim to restore.

Taken together we have shown here that repeated exposure of the peritoneal microenvironment to PD fluid instillation in mice led to a gradual change in phenotype as well as activation response of peritoneal resident cells. These changes were relatively long lasting and resulted in a more vigorous response to inflammatory triggers. Thus, targeting MΦres and preventing excessive inflammatory responses in PD patients may pose an exciting novel approach to limit PD-related pathology.

## Materials and Methods

### Ethics Statement

All animal experiments were performed in accordance with the UK Animals (Scientific Procedures) Act of 1986 under a Project License (70/8548) granted by the UK Home Office and approved by the University of Manchester Ethical Review Committee.

### Mice and in vivo treatments

Eight to 13 week old male and female C57BL/6 mice were obtained from a commercial vendor (Envigo, Hillcrest, UK). Cx3cr1^CreER^ mice [58] were obtained from the Jackson Laboratory (Bar Habor, ME) bred in-house and crossed with R26R-EYFP animals (The Jackson Laboratory, Bar Habor, ME) [81] to generate Cx3cr1^CreER^:R26-eyfp mice. All animals were maintained in groups of 4-6 animals in specific pathogen–free facilities at the University of Manchester. Experimental mice were age and sex matched and randomly allocated to treatment groups using a computer-based randomization technique and treatments were given following this a priori determined order. Euthanasia was performed by asphyxiation in carbon dioxide in a rising concentration. No experimental animals allocated to experimental groups were excluded from the analysis. Researchers were not blinded to group allocations. Animal numbers were based on initial power calculations of key determinants derived from preliminary experiments. Animals were injected i.p. with 500 μL peritoneal dialysis fluid (Physioneal 40, 3.86% glucose, Baxter HealthCare Ltd., Compton, UK) for the indicated number of injections supplemented with 40 mM Methylglyoxal (Sigma Adlrich) where indicated. Injections were carried out daily or every other day for three or five days per week for up to 4 weeks (maximum 14 injections per animal).

Control animals were left untreated or received an equal volume of PBS i.p. as indicated. For the induction of inflammatory responses PD fluid treated mice received intra-peritoneal injections of 400 μL 4% Brewer modified thioglycollate medium (BD Biosciences, San Jose, CA) or sterile saline as control. The attenuated Salmonella enterica serovar Typhimurium strain SL3261 (ΔaroA) [82] was cultured overnight at 37 °C in a shaking incubator from frozen stock in Luria-Bertani broth with 50 μg/mL streptomycin. The following morning, culture was diluted in fresh Luria-Bertani broth with 50 μg/mL streptomycin and incubated at 37 °C in a shaking incubator to ensure the bacteria were in the growth phase. CFU/mL was estimated by the OD600 reading. Animals were pre-treated with 20 mg streptomycin 1 day prior to oral infection with ∼1×10^8 CFU Salmonella Typhimurium diluted in PBS. Infectious doses and peritoneal bacterial burdens were enumerated by plating inocula or peritoneal exudate cells in 10-fold serial dilutions in PBS on LB-Agar plates.

### Cell-isolation

Peritoneal cavity exudate cells (PEC) were obtained by washing the cavity with 10 mL lavage media comprised of RPMI 1640 (Sigma-Aldrich, Dorset, UK) containing 2 mM EDTA and 1% L-Glutamine (Thermo Fisher Scientific, Waltham, MA). Erythrocytes were removed by incubating with red blood cell lysis buffer (Sigma-Aldrich). Cellular content was assessed by cell counting using Viastain AO/PI solution on a Cellometer ® Auto 2000 Cell Counter (Nexcelom Bioscience, Manchester, UK) in combination with multicolor flow cytometry.

### Flow cytometry

Equal numbers of cells were stained with Zombie UV viability assay (Biolegend, London, UK). All samples were then blocked with 5 μg/mL anti CD16/32 (93; BioLegend Cat# 101301, RRID:AB_312800) and heat-inactivated normal mouse serum (1:10, Sigma-Aldrich) in flow cytometry buffer (0.5% BSA and 2 mM EDTA in Dulbecco’s PBS) before surface staining on ice with antibodies to F4/80 (BM8; Cat# 123146, RRID:AB_2564133), SiglecF (E502440, BD Biosciences Cat# 562681, RRID:AB_2722581), Ly6C (HK1.4; Cat# 128033, RRID:AB_2562351), Ly-6G (1A8; Cat# 127612, RRID:AB_2251161), TCRβ (H57-597; Cat# 109226, RRID:AB_1027649), CD11b (M1/70; Cat# 101242, RRID:AB_2563310), CD11c (N418; Cat# 117334, RRID:AB_2562415), I-A/I-E (M5/114.15.2; Cat# 107622, RRID:AB_493727), CD19 (6D5; Cat# 115523, RRID:AB_439718), CD115 (AFS98; Cat# 135530, RRID:AB_2566525), CD73 (TY/11.8; Cat# 127206, RRID:AB_2154094), CD102 (3C4 (MIC2/4); Cat# 105604, RRID:AB_313197), Tim4 (RMT4-54; Cat# 130010, RRID:AB_2565719), CD206 (MR6F3; Cat# 141723, RRID:AB_2562445), Vsig4 (NLA14; Thermo Fisher Scientific Cat# 17-5752-82, RRID:AB_2637429), CD226 (10E5; Cat# 128816, RRID:AB_2632821), Sca1/Ly6A (D7; Cat# 108139, RRID:AB_2565957). All antibodies were purchased from Biolegend unless stated otherwise.

Detection of intracellular activation markers was performed directly ex vivo. Cells were stained for surface markers then fixed and permeabilized for at least 16 h using the eBioscience™ Foxp3 / Transcription Factor Staining Buffer Set (Thermo Fisher Scientific). Cells were then stained with directly labeled Abs to NOS2 (CXNFT; Thermo Fisher Scientific Cat# 25-5920-82, RRID:AB_2573499), Arg1 (polyclonal, R and D Systems Cat# IC5868A, RRID:AB_2810265), Ki67 (B56, BD Biosciences Cat# 563755, RRID:AB_2738406), Annexin V (Biolegend), Gata-6 (D61E4; Cell Signaling Technology Cat# 26452, RRID:AB_2798924) or purified polyclonal rabbit anti-Relmα (PeproTech Cat# 500-P214-50ug, RRID:AB_1268843) and biotinylated anti-Ym1/2 (polyclonal, R and D Systems Cat# BAF2446, RRID:AB_2260451) followed by Zenon anti–rabbit reagent (Thermo Fisher Scientific Cat# Z25302, RRID:AB_2572214) or streptavidin BUV 737 (BD Biosciences Cat# 612775, RRID:AB_2870104), respectively.

Samples were acquired on a BD LSR II or BD FACSymphony using BD FACSDiva software (BD Biosciences) and post-acquisition analysis performed using FlowJo v10 software (BD Biosciences).

### Cell-culture experiments

For in vitro stimulation of PD fluid-conditioned cells, whole PEC were counted as described above and seeded to 96-well U bottom plates at 3×10 5 cells per well in RPMI 1640 containing 5 % foetal bovine serum, 2 mM L-glutamine, 100 U/mL penicillin and 100 μg/mL streptomycin and stimulated with lipopolysaccharide (LPS, 100 ng/mL; Salmonella enterica ser. Typhimurium; Sigma-Aldrich) and recombinant murine Interferon γ (IFNγ, 20 ng/mL PeproTech EC Ltd.) or medium alone for 6 h or with murine recombinant Interleukin–4 (rIL-4, 20 ng/mL, PeproTech EC Ltd.) for 24 h and analysed for MΦ activation markers by flow cytometry.

### Apoptotic cell uptake assay

Uptake of apoptotic cells by peritoneal MΦres was assessed as previously described (52). Briefly, thymocytes were collected from naive animals by mincing thymi through 2 μm gauze until completely homogenized. Erythrocytes were removed by incubating with red blood cell lysis buffer (Sigma-Aldrich). Thymocytes were resuspended at 1×10^7 cells/mL in complete DMEM and incubated in the presence of 0.1 µM dexamethasone (Sigma-Aldrich) at 37 °C for 18 h. This produced >90 % apoptosis, as assessed by Viastain AO/PI staining measured on a Cellometer ® Auto 2000 Cell Counter (Nexcelom Bioscience). Subsequently, apoptotic thymocytes were washed twice with PBS and resuspended in PBS at 10^6 cells/mL containing 40 ng/mL pHrodo-SE (Thermo Fisher Scientific) and incubated at RT for 30 min. Thereafter the cells were washed twice with PBS and resuspended in RPMI containing 5 % foetal bovine serum, 2 mM L-glutamine, 100 U/mL penicillin and 100 μg/mL streptomycin. Unstained apoptotic cells served as staining control.

### RNA-isolation and NanoString analysis

Whole PEC were isolated as described above and total RNA isolated using Tri-reagent (Thermo Fisher Scientific) as described by the manufacturer. RNA concentration was determined using the Qbit and RNA-BR kit (Thermo Fisher Scientific). Samples were diluted and 100 ng of RNA was processed for running on a NanoString nCounter FLEX system using the nCounter Mouse Fibrosis V2 panel (NanoString Technologies Inc., Seattle, WA). Raw counts were normalized to internal spike-in controls and the expression of 10 stable housekeeping genes using the geNorm algorithm and differential gene expression calculated using the nSolver Analysis software 4.0 Advanced Analysis tool (NanoString Technologies). The datasets for this study can be found in Figshare available under DOI:10.48420/14635917.

### Statistical analysis

Statistical analysis was performed using Prism 8 for Mac OS X (v8.2.1, GraphPad Prism, RRID:SCR_002798). Differences between groups were determined by t-test or ANOVA followed by Tukey’s or Dunn’s multiple comparison-test. In some cases data was log-transformed to achieve normal distribution as determined by optical examination of residuals. Where this was not possible a Mann-Whitney or Kruskal-Wallis test was used. Percentages were subjected to arcsine transformation prior to analysis. Differences were assumed statistically significant for P values of less than 0.05.

## Author contributions

-TES: Resources, Writing – Review & Editing, Funding Acquisition

-TNS: Investigation, Resources, Writing – Review & Editing

-RL: Resources, Writing – Review and Editing, Clinical advice

-SEH: Conceptualization, Writing – Review and Editing

-DR: Conceptualization, Formal Analysis, Validation, Investigation, Writing -Original Draft Preparation, Writing – Review and Editing, Visualization, Project Administration, Funding Acquisition

## Competing interests

The authors declare that they have no competing interests.

### List of abbreviations

GDP: glucose degradation products
MΦ: macrophage
MΦres: tissue resident macrophage
MΦmono: monocyte-derived macrophage
MGO: Methylglyoxal
PEC: peritoneal exudate cells
PD: peritoneal dialysis
HD: Haemodialysis

## Acknowledgements

This work was supported by the Medical Research Council UK (MR/P02615X/1, DR) and the Medical Research Foundation / Asthma UK (MRFAUK-2015-302, TES). TNS was supported by a BBSRC Discovery Research Fellowship (BB/S01103X/1, TNS), RL was supported by a Wellcome Trust Senior Fellowship award (202860/Z/16/Z, RL) and SEH and DR were supported by an MRC Research Grant (MRC MR/S02560X/1, SEH). The nCounter transcript analysis was supported by a University of Manchester Immunology Gene Expression Panel Grant (NanoString Technologies Inc., Seattle, WA).

We thank Dr. Gareth Howell from the Flow Cytometry Core Facility at the University of Manchester for assistance with the flow cytometry analyses. We’d also like to thank Sister Trish Smith from the Royal Manchester Children’s Hospital for the provision of Physioneal 40, Dr John Grainger (University of Manchester, UK) for the provision of the Cx3cr1^CreER^ : R26-eyfp mice as well as Prof. Judith E. Allen (University of Manchester, UK) for use of her Home Office animal licence and critical appraisal of the work.

---

**Figure S1:**
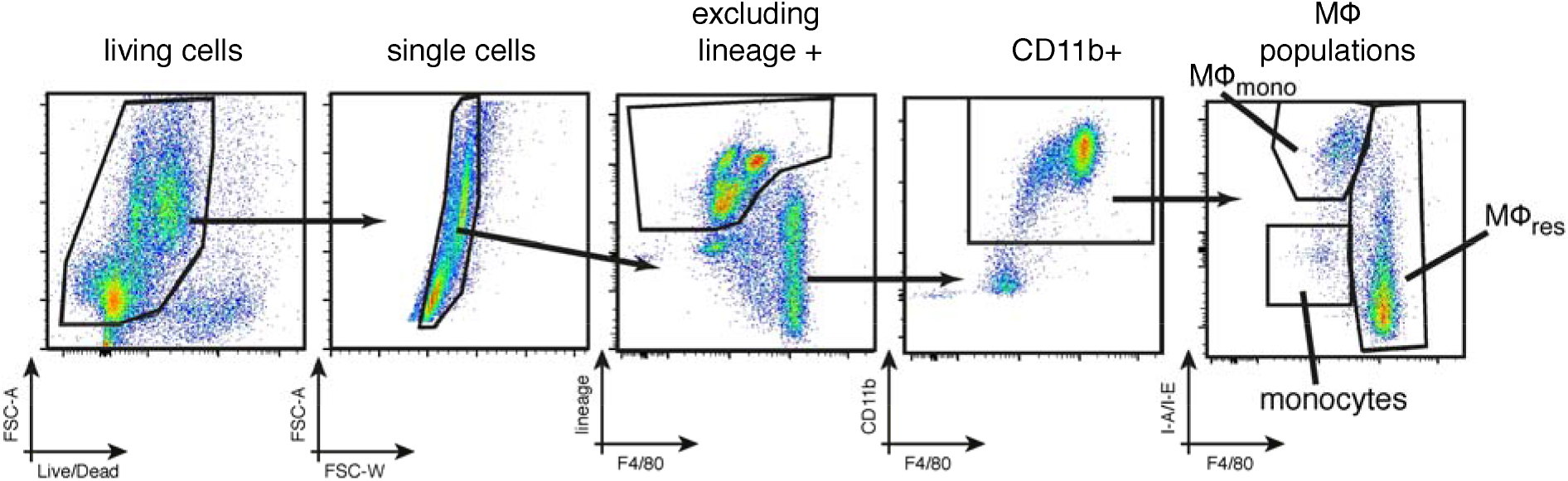
Gating strategy employed to identify MΦres, MΦmono and Ly6C high monocytes.

**Figure S2:**
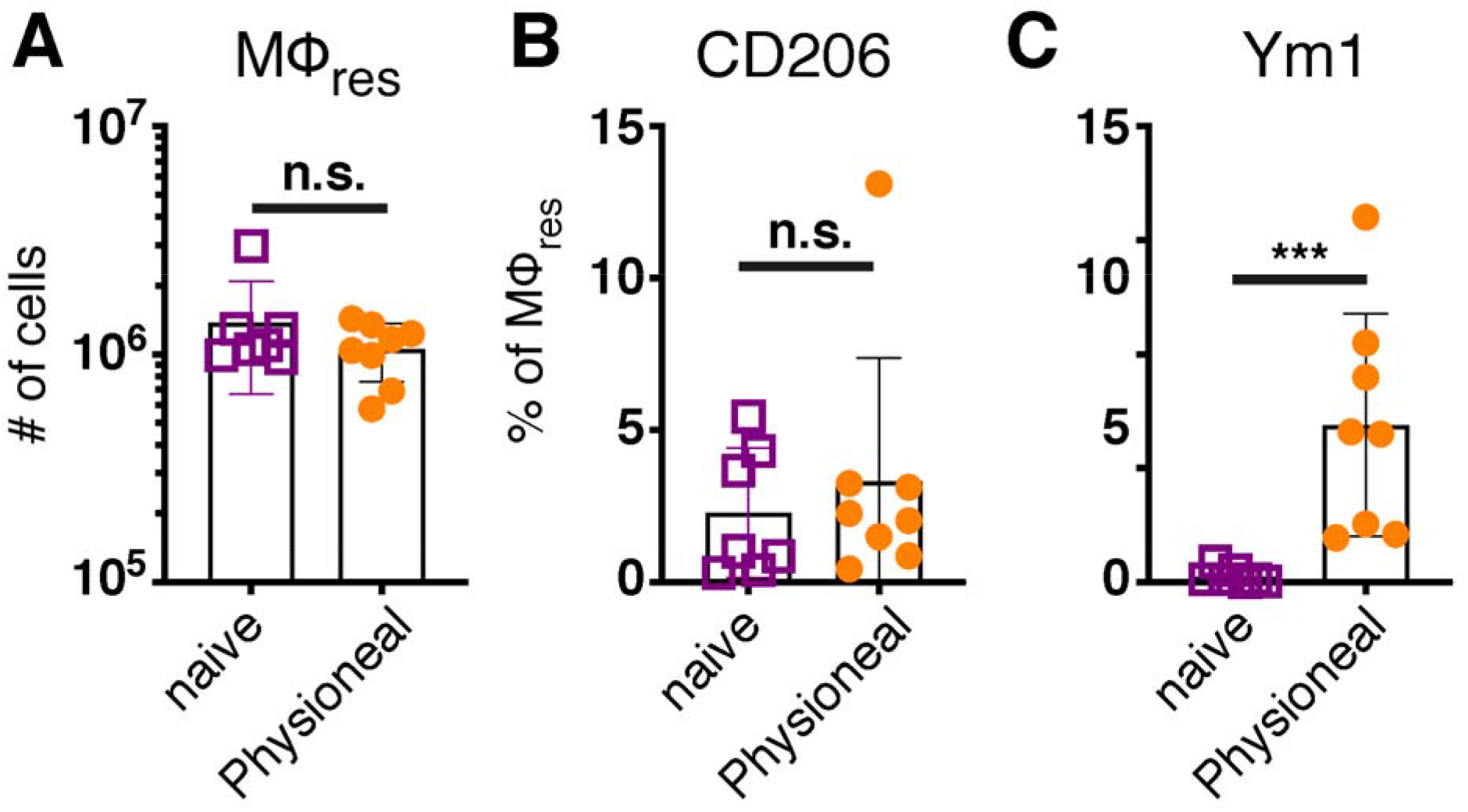
The effects of PD fluid injection on peritoneal MΦres are transient. C57BL/6 mice were injected with Physioneal (orange circle) or left untreated (purple square) and whole PEC isolated 24 h p.i.. Cells were analysed by flow cytometry to determine the number of MΦres (A) expression of cellular activation markers (B & C). Datapoints depict individual animals and bars indicate mean and SD. Data pooled from 2 independent experiments using 3-4 animals per group. Data analysed using a Mann-Whitney-U test after transformation. n.s.: not significant; ***: p<0.001

**Figure S3:**
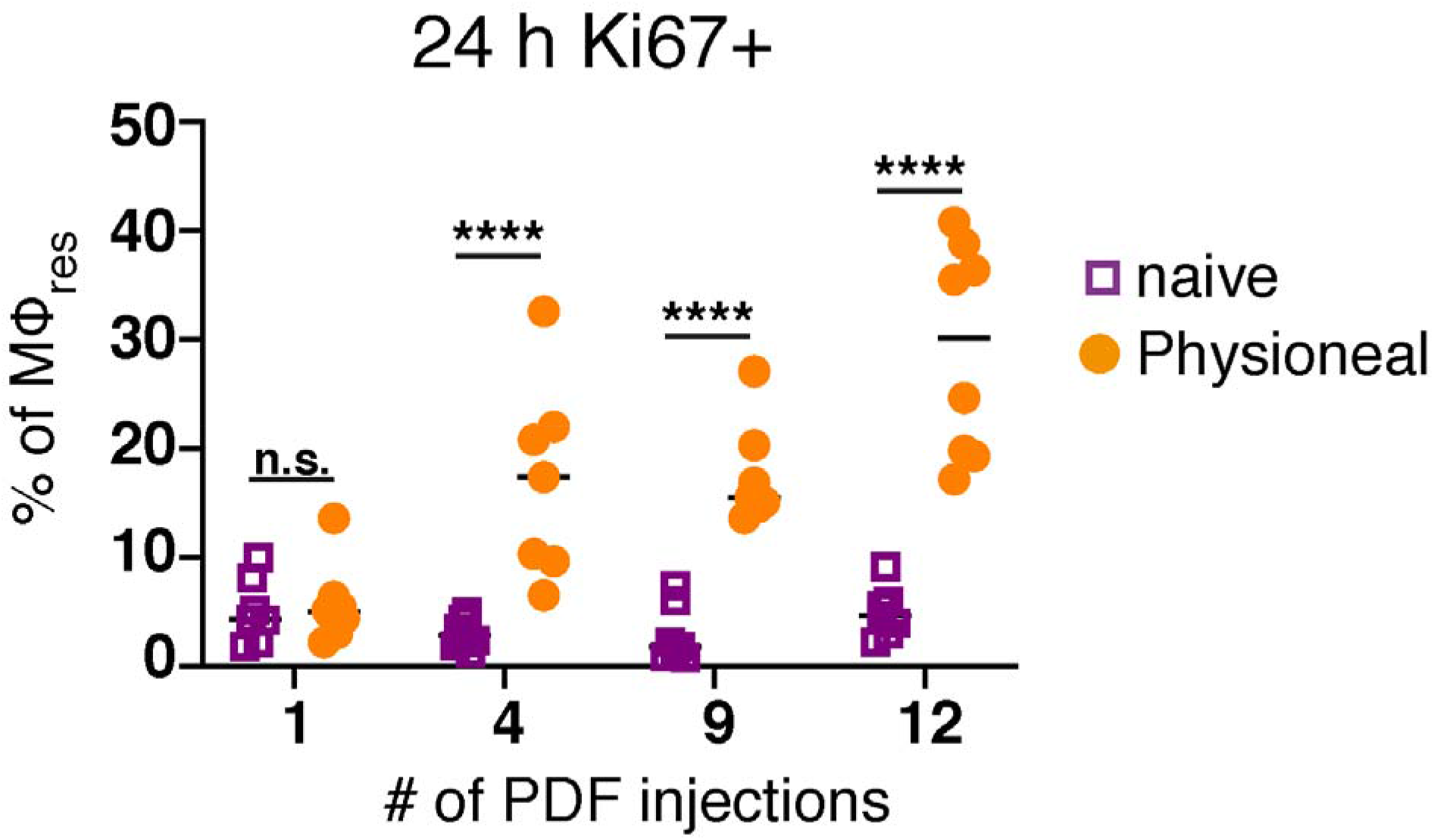
MΦres re-populate the peritoneal cavity through local proliferation following PD fluid injection. C57BL/6 mice were injected with Physioneal (orange circle) or left untreated (purple square) 5 times a week for the indicated number of injections. At each timepoint whole PEC were isolated 24 h after the last injection and analysed by flow cytometry for intracellular expression of Ki67. Datapoints depict individual animals and lines indicate mean. Data from separately performed experiments for each timepoint. Data analysed using 2-way ANOVA followed by Sidak’s multiple comparison test after transformation. n.s.: not significant; *: p<0.05; **: p<0.01; ****: p<0.0001

**Figure S4:**
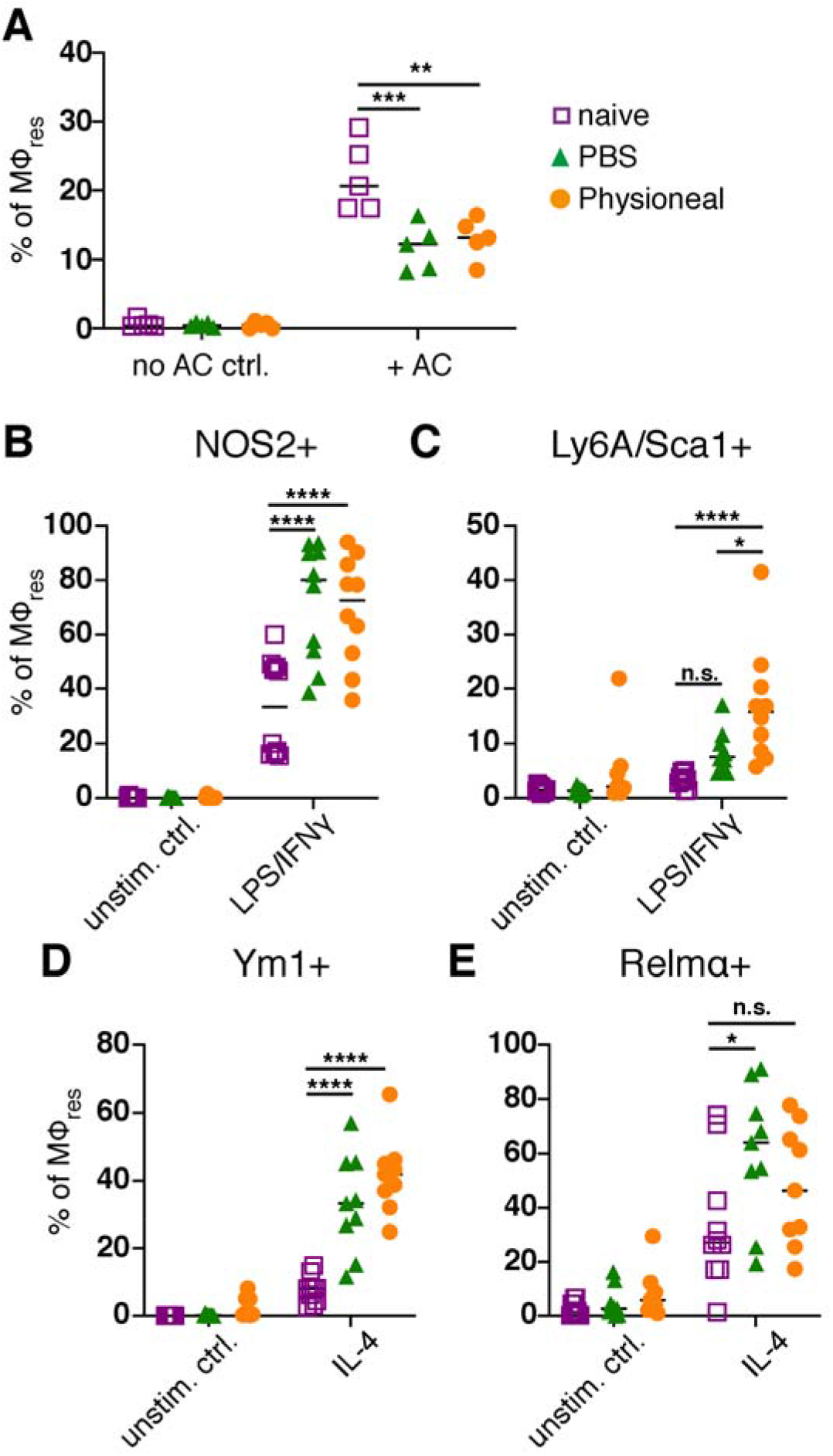
Altered MΦres phenotype is maintained after discontinuation of PD fluid injection. C57BL/6 mice were injected 5 times a week for a total of nine injections with Physioneal (orange circles), sterile PBS (green triangles) or left untreated (purple squares). 24 h after the last injection whole PEC were isolated and stimulated in vitro with (A) pHRodo labelled apoptotic cells for 90 minutes, (B) LPS/IFNγ for 6 h or (C) IL-4 for 24 h and analysed by flow cytometry. Datapoints depict individual animals and lines indicate mean. Data pooled from two independent experiments. Data analysed using 2-way ANOVA followed by Tukey’s multiple comparison test after transformation. n.s.: not significant; *: p<0.05; **: p<0.01; ****: p<0.0001; ####: P<0.0001

**Figure S5:**
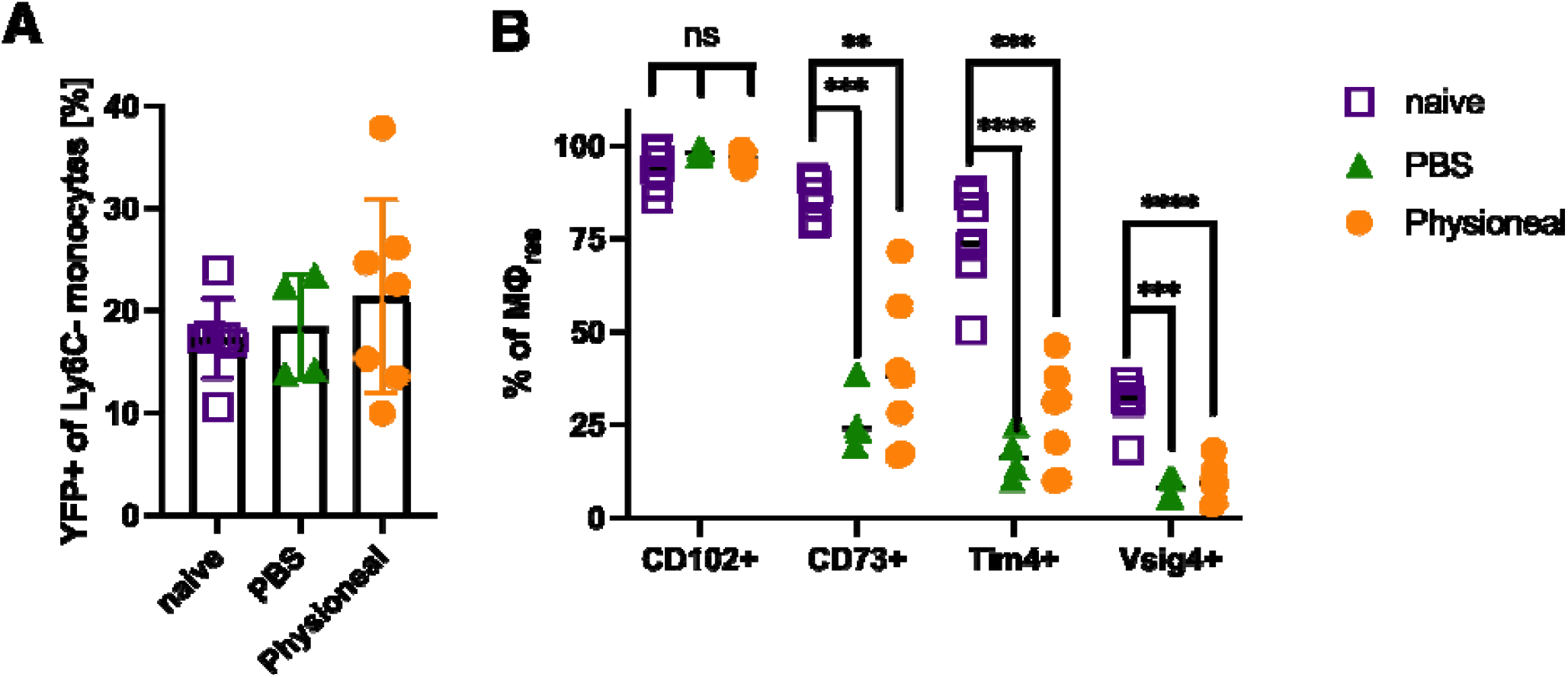
Cx3cr1^CreER^:R26-eyfp mice were injected daily with 5 mg tamoxifen for 5 consecutive days. Simultaneously, animals received daily injections, five times a week for a total of 9 injections of dialysis fluid (Physioneal, orange circles), PBS (green triangles) i.p. or were left untreated (naive, purple squares). A) Percent eYFP positive, Ly6C-monocytes (CD115+ CD19 -) in the blood. B) Expression of CD102, CD73, Tim4 and Vsig4 on peritoneal MΦ_res_ in the mice analysed in A.

